# Coated Bacterial Enzymes: A one-step approach for enzymatic purification and immobilization

**DOI:** 10.64898/2026.07.08.735634

**Authors:** Andrea C. Ramírez, Inés Harguindeguy, Melisa S. Homse, Alan E. Sabetta, Sebastián F. Cavalitto, Gastón E. Ortiz

**Author notes:** These authors contributed equally (Andrea C. Ramírez, Inés Harguindeguy). Corresponding author: Prof. Dr. Gastón E. Ortiz.

## Abstract

The purification of industrial enzymes typically relies on costly, multi-step chromatographic protocols. To address this, we developed a novel platform termed Coated Bacterial Enzymes (CBEs), which enables one-step purification and immobilization of recombinant proteins fused to the SlpA cell wall binding domain. As a proof of concept, we used a β-galactosidase from *Bifidobacterium bifidum* of dairy relevance. The chimeric enzyme BbgII-SlpA was expressed in *Escherichia coli* and captured from crude lysate onto glutaraldehyde-inactivated *Bacillus subtilis* cells via SlpA domain. Binding was characterized by a dissociation constant (K_d_) of 16.2 μM and maximum binding capacity (B_max_) of 144 μmol/g. The resulting CBE biocatalyst exhibited optimal activity at pH 6.0 for ONPG and lactose, with a broader pH profile than the free enzyme. Optimal temperatures were 60 °C for ONPG and 50 °C for lactose, and CBE retained >80% activity after 390 min at 45 °C, compared to 20% for the free enzyme. Catalytic efficiencies (kcat/Km) were 2.62 × 10^6^ M^−1^·s^−1^ for ONPG and 4.40 × 10^2^ M^−1^·s^−1^ for lactose. Moreover, CBE showed improved tolerance to cations such as Ca^2+^ and Fe^2+^. These results suggest that the CBE platform offers a cost-effective alternative for producing high-purity, immobilized enzymes for diverse industrial bioprocesses.

## 1. Introduction

Lactose is a disaccharide composed of glucose and galactose present in all mammalian milks, including cow, goat, sheep, and human milk. Its concentration ranges from approximately 4.7% to 7% (w/v), and its primary function is to provide the infant with a source of carbon and energy [1]. This disaccharide is hydrolyzed into glucose and galactose in the small intestine of mammals during the neonatal period by the enzyme β-galactosidase (EC 3.2.1.23), commonly known as lactase [2]. In adult mammals, the specific activity of this enzyme declines, leading to difficulties in lactose assimilation [3]. Unabsorbed lactose continues its transit to the large intestine, where it is fermented by the gut microbiota, producing H_2_, CO_2_, CH_4_, and short-chain fatty acids. These products cause the characteristic symptoms of lactose intolerance, including abdominal pain and bloating, diarrheic, and flatulence [1,3]. If these symptoms persist, they can lead to deterioration of the intestinal mucosa, which may contribute to the development of more severe pathologies such as celiac disease, infectious enteritis, or Crohn’s disease [3]. According to the literature, 70% of the world population is estimated to be predisposed to some degree of lactose intolerance [4].

Due to the impact of this syndrome on public health and nutrition, numerous strategies have been developed to produce lactose-reduced milk (LRM). These strategies involve chemical, physical, or enzymatic treatment of milk, with enzymatic methods being the most widely adopted [5][6]. Industrial production of LRM via the enzymatic method is typically carried out after milk pasteurization and involves treating pasteurized milk with the catalyst (lactase), either free or immobilized on an inert matrix (formulated catalyst), for 24 hours at 8 °C [7]. After catalysis, the LRM is ultrapasteurized using ultra-high temperature (UHT) treatment, which consists of heating the LRM to 150 °C for a few seconds. This process inactivates the catalyst and significantly reduces the microbial load [8].

The LRM process requires a catalyst free of contaminating enzymes, such as proteases. The presence of proteases leads to LRM with undesirable organoleptic properties such as bitterness, or increases free amino acids and small peptides that favour the formation of Maillard reaction products, some of which are undesirable (e.g., acrylates) [7]. Therefore, lactase must be obtained with a high degree of purity during its production process to be used in LRM manufacturing. Other characteristics required for lactase include good residual activity at temperatures close to those used during the hydrolysis process (8 °C), as well as good activity and stability at pH values of 6–7 and at concentrations of approximately 0.7–1.5% (w/v) for K^+^, Mg^2+^, Ca^2+^, and Na^+^, the main cations present in milk [9–12]. Lactases from different microorganisms such as *Kluyveromyces lactis*, *Kluyveromyces fragilis*, *Bacillus subtilis*, and *Bifidobacterium bifidum* have been used for this type of process because they exhibit optimal activity at pH 6–7. However, their optimal temperatures are around 37 °C for the genus *Kluyveromyces* and 50 °C for *Bacillus* and *Bifidobacterium* [13].

Due to these enzyme properties, several companies supply lactase enzymes, including two neutral lactases (pH 6–7) from *K. lactis* (Lactozym®) and recombinant BbgII from *B. bifidum* (Saphera®) [11,14]. BbgII belongs to glycosyl hydrolase family 42 and prefers β-D-(1→6)-linked galactosides over lactose, with an optimal activity at pH 6.5 and 40–50 °C [11,14].

As mentioned above, the production of LRM requires a high degree of enzyme purity. Most purification protocols use two or more column fractionation steps and involve techniques such as gel filtration, ion exchange, and affinity chromatography. Multi-step purification is time-consuming, expensive, and generally results in low yields [15,16]. To overcome this limitation, one-step purification methods have been employed. These methods involve cloning the gene of interest fused to a polypeptide tag that allows affinity purification between the tag and a specific ligand, such as the use of polyhistidine tags and Ni^2+^ affinity chromatography [17]. However, these affinity purification techniques are usually expensive because they require the preparation of specific affinity matrices.

To address this issue, research has focused on the search for new tags and low-cost ligands. In this context, purification systems based on the interaction of chitin-binding, cellulose-binding, or starch-binding domains with their respective matrices formulated with chitin, cellulose, or starch have been reported [18,19]. For instance, Gennari et al. achieved one-step purification and immobilization of a β-galactosidase from *Kluyveromyces* sp. using a cellulose-binding domain (CBD) tag on magnetic cellulose supports, enabling rapid enzyme recovery and enhanced thermal stability [20]. Other affinity strategies include the use of polyhydroxyalkanoate-binding domains, chitin-binding modules, and carbohydrate-binding modules, as comprehensively reviewed by Duan et al. [21].

Recently, Muruaga et al. developed a novel molecular tagging method based on the cell wall-binding domain (SlpA^284−444^) of the S-layer protein A from *Lactobacillus acidophilus*. This tag enables the affinity purification of recombinant proteins using a low-cost matrix derived from *B. subtilis*. Muruaga and colleagues successfully validated this purification method using GFP protein, with yields and purity comparable to commercial immobilized metal affinity chromatography (IMAC) [22].

On this basis, in this work we have developed a novel biocatalyst platform that we have termed Coated Bacterial Enzymes (CBEs). This approach employs inactivated *Bacillus subtilis*, which retains its structural integrity and provides a low-cost biological carrier, to simultaneously purify and immobilize a recombinant β-galactosidase fused to the SlpA cell wall-binding domain. Accordingly, the present study aims to expand the scope of this SlpA-based technology toward enzymatic applications by evaluating the system using the dairy industry model enzyme BbgII from *Bifidobacterium bifidum*. We report the optimization of BbgII-SlpA expression, its binding to the *B. subtilis* carrier, and the functional characterization of the resulting CBE biocatalyst (pH and temperature optima, thermal stability, kinetics, and cation sensitivity). Our results indicate that the CBE system offers a straightforward, low-cost alternative for obtaining high-purity, immobilized β-galactosidase, with potential application in the production of lactose-reduced milk.

## 2. Materials and Methods

### 2.1. Bacterial strains and culture conditions

*Escherichia coli* BL21-CodonPlus (DE3)-RIPL strain (*E. coli* BL21) was used as the host for heterologous protein expression. *Bacillus subtilis* var. natto (ATCC 15245) was used as the antigen-presenting carrier. *E. coli* and *B. subtilis* strains were grown in Luria-Bertani broth (LB, pH 7.0) in a shaking incubator (200 rpm) or on LB-agar plates at 37°C. Bacterial cell growth was monitored by measuring the absorbance at 600 nm (OD_600_) using a spectrophotometer [23]. For *E. coli* BL21 transformant selection, kanamycin (Sigma-Aldrich) was used at a final concentration of 50 μg/ml on LB or LB agar medium. For *B. subtilis* dry cell weight (DCW) was calculated by multiplying the OD_600_ value by 0.322 g/OD_600_ conversion factor [24].

### 2.2. Expression vector, enzyme design and E. coli transformation

To generate the expression vector pET-BbgII-SlpA, the SlpA sequence comprising amino acids 310 to 470 of the carboxyterminal fragment of the SlpA protein from *L. helveticus* (GenBank ID: MBW7998749.1), and the BbgII amino acids sequence (GenBank ID: WP_270548944.1) were subjected to optimization for *E. coli* codon usage using the J-Cat software (https://www.jcat.de). The resultant optimized DNA sequence, BbgII-SlpA, was chemically synthesized by General Biosystems, Inc. and subsequently cloned into the pET-29c(+) vector to give the expression plasmid (pET-BbgII-SlpA). One hundred ng of the resulting expression plasmid were transformed into chemically competent *E. coli* BL21 cells by heat shock protocol, and positive transformants were selected on LB-kanamycin agar plates, as described by Sambrook and Russell [25]. After selection, the recombinant *E. coli* colonies were screened using the small-scale protocol for induction of heterologous protein expression [26].

### 2.3. Optimization of BbgII-SlpA Expression

For optimization of soluble BbgII-SlpA, a one-factor-at-a-time approach was performed. Recombinant *E. coli BL21* cultures were grown as described in section 2.1 to an OD_600_ of ~0.5. Induction temperature (22, 28, or 37 °C) was evaluated using 1.0 mM IPTG for 16 h. IPTG concentration (1.0, 0.5, 0.2, or 0.1 mM) was tested at 22 °C for 16 h. Induction time (0–8 h, with a 16 h endpoint) was evaluated at 22 °C using 0.1 mM IPTG. After each condition, cells were lysed and fractionated as described in section 2.4, and soluble/insoluble protein yields were quantified by SDS-PAGE and densitometry (section 2.5).

### 2.4. Production of BbgII-SlpA and cell lysis

For recombinant BbgII-SlpA production, *E. coli* BL21 cultures were grown as described in section 2.1 to reach an OD_600_ value of ~0.5. Cultures were then cooled to 22 °C, and induced with 0.1 mM IPTG (Promega) for 16 h at 22 °C. Cells were harvested by centrifugation at 5,000 × g for 20 min at 4 °C and resuspended in lysis buffer (50 mM Tris-HCl, pH 7.5; 150 mM NaCl; 1 mM EDTA; 0.3 mg/mL lysozyme). The suspension was incubated for 30 min at 37 °C with gentle agitation. Cells were disrupted by sonication (2’’ x 30 cycles at 30% amplitude) using an ultrasonic homogenizer equipped with a No. 6 tip (SCIENTZ IID model JY96-II). The homogenate was centrifuged at 32,000 × g for 30 min at 4 °C to separate the soluble (supernatant) and insoluble (pellet) fractions. The supernatant, soluble fraction, was collected and 40 μL of sample was taken for analysis by SDS-PAGE. Pellet was washed three times with wash buffer (5 mM Na_2_HPO_4_, 150 mM NaCl, 5 mM KH_2_PO_4_, pH 7.4, 2% Triton™ X-100, 1 M urea), and then solubilized in solubilization buffer (5 mM Na_2_HPO_4_, 150 mM NaCl, 5 mM KH_2_PO_4_, pH 7.4, 8 M urea). Soluble proteins from inclusion bodies were recovered from the supernatant after centrifugation at 32,000g for 30 min at 4°C, and 40 μL of sample was used for analysis by SDS-PAGE.

### 2.5. SDS-PAGE and densitometry analysis

The soluble, insoluble, and total protein fractions were diluted in a ratio of 4:1 in 5X loading buffer (0.25 M Tris-HCl, pH 6.8, 50% glycerol, 10% SDS, 0.1% bromophenol blue, 5% β-mercaptoethanol). The samples were heated at 90°C for 10 min and 20 µL of each were loaded in a 15% SDS-PAGE gel. The electrophoresis was performed at 150 V for 60 min in a Mini-Protean Tetra cell (Bio-Rad) and gels were stained with Coomassie Brilliant Blue dye according to Sambrook and Russel [25]. For BbgII-SlpA quantification, the SDS-PAGE gels were loaded with 5; 10 and 15 µL of 0.1 µg/µL bovine serum albumin (BSA) solution as calibration curve. After staining, the gels were digitized using a scanner Epson Stylus CX3700 and the images were analyzed by ImageJ densitometry software [27]. The percentage of soluble or insoluble BbgII-SlpA was calculated as (amount of protein in the respective fraction / total amount of protein) × 100.

### 2.6. Preparation of bacterial carrier

The coated bacterial enzyme (CBE) was prepared using chemically inactivated *B. subtilis* as the bacterial carrier. Briefly, an overnight culture of *B. subtilis* was harvested by centrifugation at 5,000 × g for 10 min at 4 °C. The bacterial pellet was washed three times with phosphate-buffered saline (PBS, pH 7.4) and inactivated by resuspension in 2.5% (v/v) glutaraldehyde overnight at 4 °C with gentle stirring. Inactivated cells were washed five times with PBS and resuspended in the same buffer to a concentration of 50 OD_600_/mL.

### 2.7. Production of CBEs

For the simultaneous purification and immobilization of BbgII-SlpA, 20 OD_600_ units of *E. coli* lysate containing soluble BbgII-SlpA (obtained as described in section 2.3) were mixed with 10 OD_600_ units of the bacterial carrier in a final volume of 500 μL. The mixture was incubated for 1 h at room temperature on a rotator. After incubation, the bacterial carrier was centrifuged at 5,000 rpm for 3 min, and the supernatant (unbound fraction) was collected and reserved for SDS-PAGE analysis. The pellet was washed three times with one volume of PBS, and an aliquot was taken to determine the amount of BbgII-SlpA bound to *B. subtilis*, expressed as protein amount per OD_600_ unit of the bacterial carrier. Protein quantification was performed by SDS-PAGE followed by densitometry analysis, as described in section 2.4.

### 2.8. Purification of recombinant BbgII-SlpA

The immobilized enzyme was liberated by three consecutive elutions with buffer pH 6.0, prepared by mixing one part of 400 mM carbonate-bicarbonate buffer pH 9.0 with one part of 1 M citrate-phosphate buffer pH 6.0. The eluate was dialyzed using a cellulose tubing (cut-off value 12.4 kDa, Sigma-Aldrich) against PBS overnight at 4°C and then stored at 4°C until further use. The protein concentration was determined using the Pierce BCA Protein Assay Kit (Thermo Fisher) and BbgII-SlpA integrity was analyzed by SDS-PAGE.

### 2.9. Determination of binding kinetic constants

The maximum binding capacity (B_max_) and the apparent dissociation constant (K_d_) were determined by a saturation assay as described by Harguindeguy et al. [28]. A fixed number of bacterial carrier cells (1×10^8^ cells) was incubated with increasing amounts of purified BbgII-SlpA (5.4–32.1 μg) in a final volume of 200 μL PBS for 30 min at 25 °C. After incubation, the cells were pelleted and washed three times with PBS, and the amount of bound protein was evaluated by SDS-PAGE and densitometry analysis as described in section 2.4. The dissociation constant (K_d_) and the maximum amount of bound protein (B_max_) were calculated using a sigmoidal model (Eq. 1).

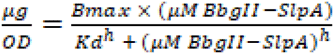

**Eq. 1.** Equation used to calculate the regression curve corresponding to the equilibrium isotherm and determine the binding parameters (B_max_ and K_d_), where B_max_ is the maximum amount (μg) of protein bound per OD of *B. subtilis*; [μM_BbgII-SlpA_] is the molar concentration of protein in the system, K_d_ (μM) is the dissociation equilibrium constant and *h* is a regression coefficient used to describe the slope of a sigmoidal curve between these two plateaus, commonly known as Hill slope.

### 2.10. Immunofluorescence microscopy

Immunofluorescence microscopy was performed as described by Harguindeguy et al. [28]. Briefly, CBEs were adhered to poly-L-lysine-coated coverslips, fixed, quenched, and blocked. Samples were then incubated with anti-SlpA mouse serum previously obtained in our laboratory [31], followed by Alexa 488 conjugated goat anti-mouse IgG antibody (Thermo Fisher). After each incubation, coverslips were washed with PBS. Finally, samples were analyzed using an epifluorescence microscope.

### 2.11. Determination of BbgII-SlpA activity

β-Galactosidase activity was assayed using, o-nitrophenyl-β-D-galactopyranoside (ONPG) and lactose. For ONPG assays, 10 μL of enzyme were added to 490 μL of 15 mM ONPG in 100 mM Na_2_HPO_4_-citric acid buffer (pH 6.0) at 37 °C for 1 min. The reaction was stopped with an equal volume of 1 M Na_2_CO_3_, and hydrolysis was measured at 420 nm (ε_420_ = 4.5 mM^−1^·cm^−1^). One unit (U) was defined as the amount of enzyme releasing 1 μmol of o-nitrophenol per minute. For lactose assays, 20 μL of enzyme were mixed with 980 μL of 140 mM lactose in the same buffer and incubated at 40 °C for 20 min. The reaction was stopped by heat inactivation (90 °C, 5 min). Released glucose was measured using a glucose enzymatic kit at 505 nm (Winner Lab^®^). One unit was defined as the amount of enzyme releasing 1 μmol of glucose per minute. Unless otherwise stated, all biochemical characterization assays were performed using 25 U of enzyme (free or immobilized) per reaction.

### 2.12. Effect of pH and temperature on the enzyme activity

The optimal pH and temperature for free and immobilized BbgII-SlpA were determined under the standard assay conditions described in section 2.11. For pH optimization, activity was measured at 37 °C in a pH range of 4.0–8.0 using 100 mM Na_2_HPO_4_-citric acid buffer, with 2 mM ONPG or 140 mM lactose as substrates. For temperature optimization, reactions were performed at 4–70 °C at the optimal pH (6.0) using the same substrates. All experiments were performed in triplicate, and the error was expressed as the standard deviation.

### 2.13. Thermal stability

Thermostability was evaluated by preincubating CBE or free enzyme at 45 °C in 100 mM Na_2_HPO_4_-citric acid buffer (pH 6.0) for various time intervals (0, 15, 30, 90, 150, 210, 270, 330, 390 min). Residual activity was measured with ONPG and lactose under the standard assay conditions (section 2.11). Inactivation constants (k_i_) were calculated using first-order rate equations, and half-lives (t_1/2_) were derived from these values.

### 2.14. Effect of metal ions

The effect of various cations (Na^+^, K^+^, Mg^2+^, Mn^2+^, Ca^2+^, Ba^2+^, Cu^+^, Zn^2+^, Co^2+^, Fe^2+^/^3+^) on CBE activity was examined. The corresponding chloride salts were added to the substrate solution (lactose 5% w/v in 20 mM sodium phosphate buffer, pH 6.0) at final concentrations of 1, 10, and 100 mM. Activity was measured as described in section 2.11. Results were expressed as relative activity (%) compared to the enzyme without added salts. All experiments were performed in triplicate.

### 2.15. Determination of kinetics constants

Kinetic parameters of the immobilized enzyme were determined for both lactose and ONPG hydrolysis. Reactions were performed under conditions described in section 2.11, using ONPG (0–20 mM) or lactose (0–200 mM) in 0.1 M sodium phosphate buffer (pH 6.0) at 37 °C. The reaction was initiated by adding 5 μg of enzyme sample and incubated for 20 min (lactose) or 1 min (ONPG). Released o-nitrophenol was quantified as described in section 2.11, and glucose was measured using a glucose enzymatic kit. Kinetic constants (K_m_, V_max_, k_cat_) were obtained by non-linear fitting of the Michaelis-Menten equation to the initial velocity data using a custom Python script (Google Colab, SciPy curve_fit). The Michaelis-Menten fit provided K_m_ (mM) and V_max_ (U/mg) with standard errors. A Lineweaver-Burk linearization (1/v vs. 1/[*S*]) was used as a complementary check. The turnover number k_cat_ (s^−1^) was calculated as k_cat_=(V_max_·MW)/60000 using the enzyme’s molecular weight MW (g/mol). Catalytic efficiency (k_cat_/K_m_) was expressed in M^−1^·s^−1^, and errors were propagated quadratically. All plots (Michaelis-Menten and Lineweaver-Burk) were generated with Matplotlib.

## 3. Results

### 3.1. Optimization of BbgII-SlpA expression in E. coli

The obtention of a soluble and properly folded recombinant enzyme is a critical step for its subsequent immobilization on bacterial carriers. One of the main factors affecting protein solubility in *E. coli* is the growth temperature during the induction and protein production phase [29]. Lowering the temperature during induction (15–25 °C) is regarded as a common approach to enhance the solubility of recombinant proteins by reducing aggregation and the formation of inclusion bodies [30][31]. Therefore, the production of BbgII-SlpA was conducted at different induction temperatures (22 °C, 28 °C and 37 °C). For each temperature, the protein expression was normalized to culture density (OD_600_), and the total amount of soluble antigen was compared. As shown in Fig. 1A, increasing the induction temperature from 22 °C to 28 °C and 37 °C caused a progressive increase in insoluble protein yield (Yield IP) and a progressive decrease in soluble protein yield (Yield SP), while the soluble/insoluble ratio (SP/IP) also decreased. The highest proportion of soluble protein was obtained at 22°C, with solubility 2.3-fold and 5.4-fold higher than at 28 °C and 37 °C, respectively.

**Figure 1.**
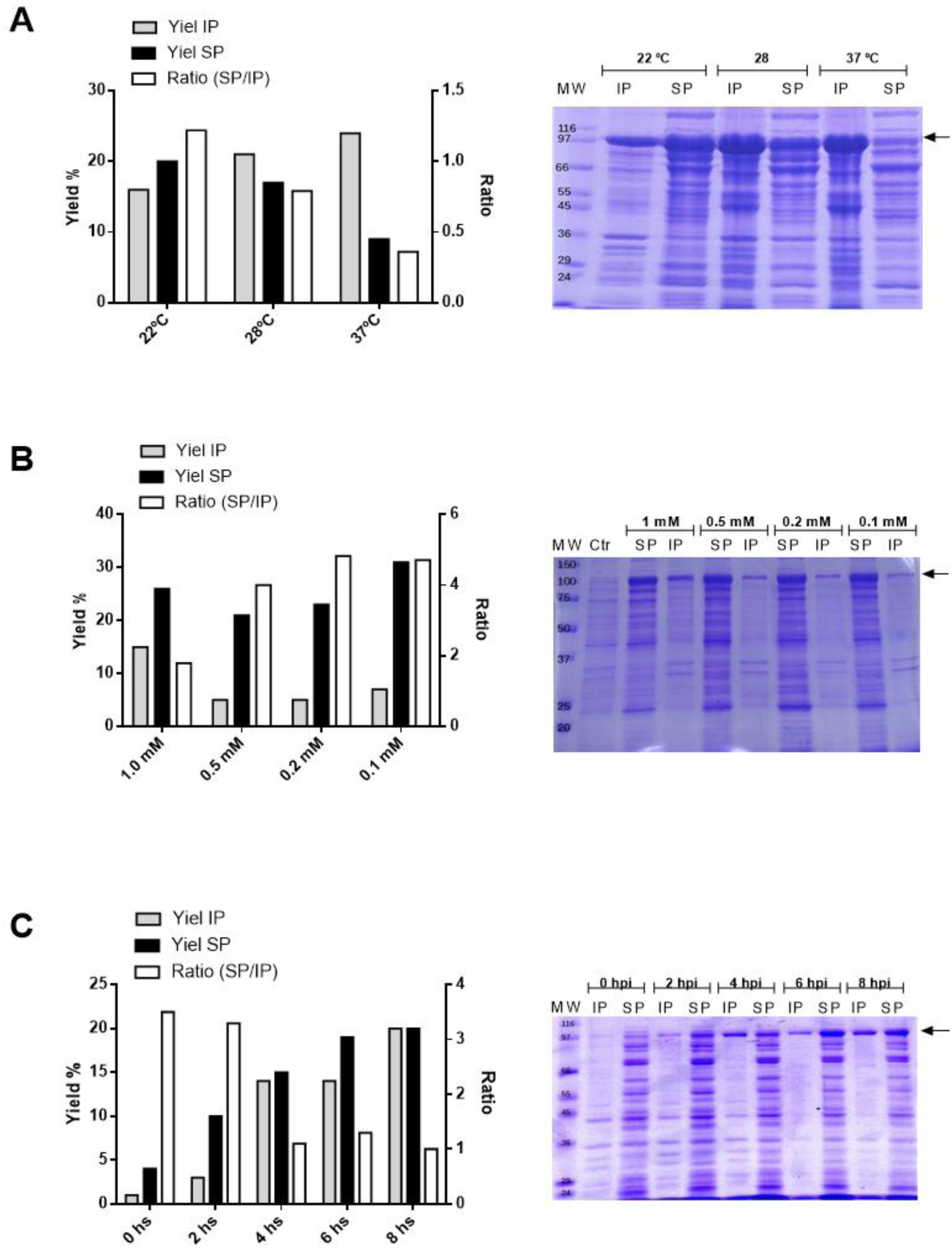
Optimization of BbgII-SlpA expression in *E. coli* BL21. Polyacrylamide gels under denaturing conditions showing cellular fractions of lysed recombinant *E. coli* subjected to different protein expression induction conditions, evaluating one factor at a time. (A) Induction temperature: cultures were induced with 1 mM IPTG and incubated for 16 h at the indicated temperatures (22, 28, or 37 °C). (B) IPTG concentration: cultures were induced with the indicated IPTG concentrations (1.0, 0.5, 0.2, or 0.1 mM) and incubated for 16 h at 22 °C. (C) Induction kinetics: cultures were induced with 0.1 mM IPTG at 22 °C, and samples were collected at the indicated times (0, 2, 4, 6, 8 h). Wells labelled (IP) indicate the insoluble protein, wells labelled (SP) indicate the soluble protein, and (Ctr) in panel B indicates the uninduced *E. coli* extract.

In addition to temperature, the inducer (IPTG) concentration and induction time were optimized using a one-factor-at-a-time approach. As shown in Fig. 1B, different IPTG concentrations (1.0, 0.5, 0.2, and 0.1 mM) were tested at 22 °C. Decreasing the inducer concentration from 1.0 mM to 0.1 mM resulted in an increase in soluble protein yield and in the SP/IP ratio. The lowest concentrations (0.2 and 0.1 mM) gave the best solubility levels and 0.1 mM IPTG was selected as the optimal concentration for subsequent experiments.

The time course of expression was evaluated at 22 °C using 0.1 mM IPTG. As shown in Fig. 1C, during the first 6 h both soluble and insoluble protein yields increased gradually. After 6 h, the soluble yield increased up to 8 h, with the SP/IP ratio remaining constant. No further improvements were observed beyond 8 h, as evidenced by the 16 h time point shown in panel Fig. 1B (22 °C, 0.1 mM IPTG), where the soluble yield did not increase.

Based on these results, the following conditions were established for the production of BbgII-SlpA. Recombinant *E. coli* BL21 was grown at 37 °C to an OD_600_ of ~0.5, then cooled to 22 °C, induced with 0.1 mM IPTG, and incubated for 16 h at 22 °C. Under these conditions, approximately 70% of the expressed BbgII-SlpA was recovered in the soluble fraction, providing sufficient protein for immobilization and purification assays. The overexpressed recombinant enzyme was confirmed by an intense band near 97–100 kDa when compared to the Sigma molecular weight marker (Fig. 1D). The calculated molecular mass of the BbgII-SlpA fusion protein (695 amino acids) is 98 kDa, in agreement with the observed band.

### 3.2. Coated bacterial enzymes production and enzyme binding properties

Once the expression conditions for BbgII-SlpA were defined, CBEs were produced as described in Materials and Methods. Briefly, *B. subtilis* var. natto cells were cultured, chemically inactivated with glutaraldehyde to generate the bacterial carrier, and then incubated with the soluble fraction of *E. coli* lysate containing BbgII-SlpA. This allowed the SlpA domain to bind specifically to the *B. subtilis* cell wall (Fig. 2A). After incubation, the bacteria were washed to remove unbound proteins, and the bound BbgII-SlpA was either used directly as CBEs or eluted for use as purified enzyme.

**Figure 2.**
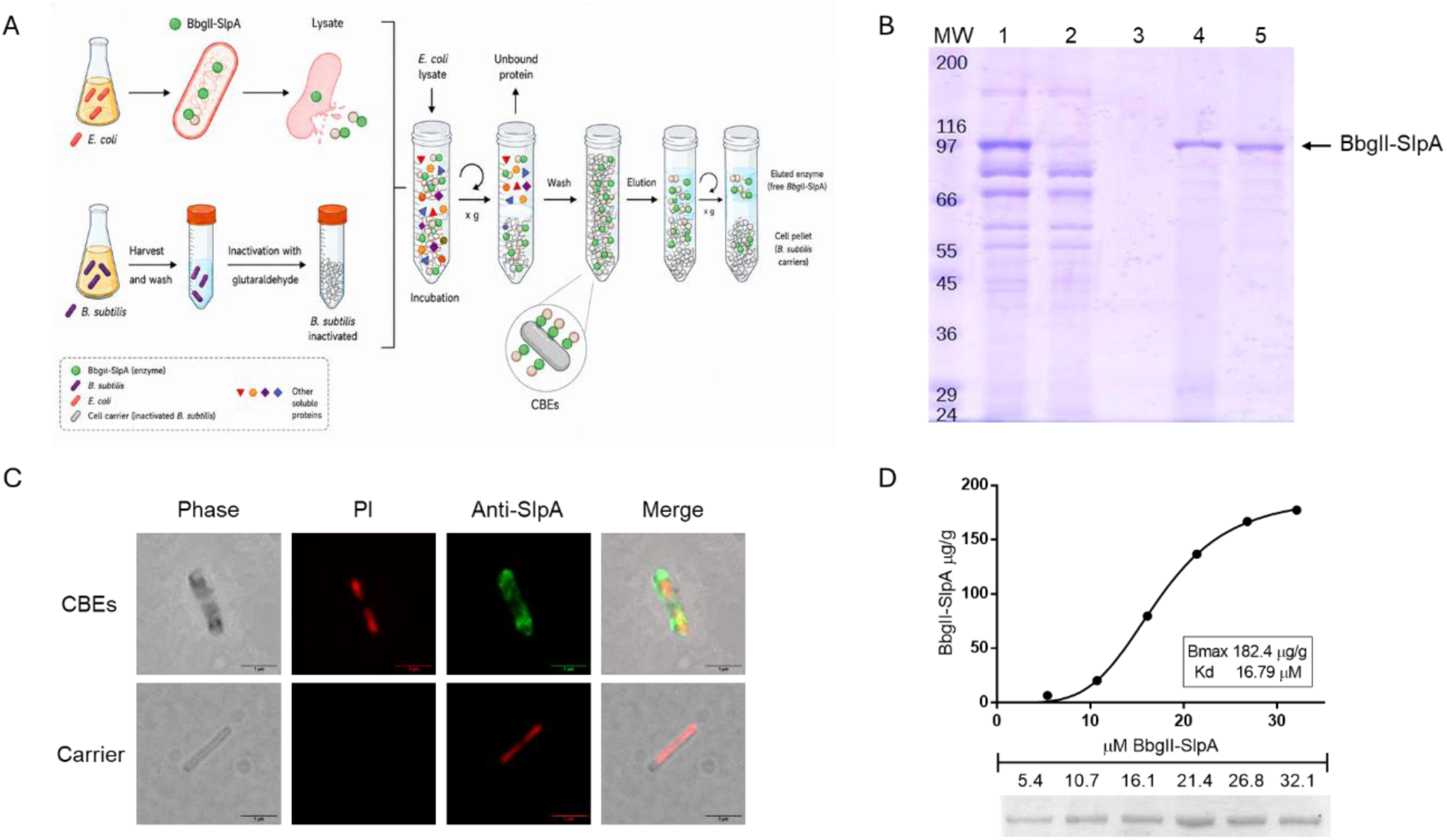
BbgII-SlpA binding to *B. subtilis*. (A) Schematic illustration of the CBEs production process: binding of BbgII-SlpA from *E. coli* lysate to inactivated bacterial carrier, followed by washing and optional elution. (B) SDS-PAGE analysis showing the efficient binding and purification of BbgII-SlpA (98 kDa band, arrow) from input (lane 1), unbound (lane 2), wash (lane 3), CBE (lane 4), and eluate (lane 5). (C) Immunofluorescence microscopy of CBE: phase contrast, propidium iodide (red, bacteria), anti-SlpA antibody (green, BbgII-SlpA), and merge. Uniform surface coating of the enzyme is observed. Scale bar = 1 μm. (D) Saturation binding isotherm obtained using increasing concentrations of purified BbgII-SlpA incubated with a fixed number of inactivated bacterial carrier (1×10^8^ cells/ml). Bound protein was quantified by densitometry from SDS-PAGE. The dissociation constant (K_d_) and maximum binding capacity (B_max_) were determined by nonlinear regression.

The efficiency of binding and purification was evaluated by SDS-PAGE (Fig. 2B). The input lysate (lane 1) contained a prominent band at approximately 98 kDa corresponding to BbgII-SlpA. After incubation with the bacterial carrier, the unbound (lane 2) showed a marked reduction of this band, indicating efficient binding. No significant protein was detected in the final wash (lane 3), demonstrating that non-specifically bound proteins were effectively removed. The CBE preparation (lane 4) clearly retained the 98 kDa band. Subsequent elution (lane 5) yielded highly purified BbgII-SlpA with minimal contaminants, confirming that the SlpA domain enables one-step purification and immobilization.

To quantify the interaction, a saturation binding assay was performed. Increasing concentrations of purified BbgII-SlpA were incubated with bacterial carrier. The amount of bound protein was determined by SDS-PAGE and densitometry. Plotting bound protein versus free protein concentration yielded a typical saturation isotherm (Fig. 2C). Non-linear regression analysis gave a dissociation constant (K_d_) of 16.2 μM and a maximum binding capacity (B_max_) of 144 μmol per gram of dry cell weight (μmol/g). These values indicate a stable and specific interaction, comparable to those reported for other proteins fuse to S-layer protein binding domains [28,32].

Finally, the localization of BbgII-SlpA on the bacterial surface was examined by immunofluorescence microscopy (Fig. 2C). CBE particles were labeled with an anti-SlpA antibody (green) and counterstained with propidium iodide (PI, red) to visualize the bacterial carrier. Phase-contrast images confirmed the presence of intact bacteria. The merge panel shows that the BbgII-SlpA signal (green) is uniformly distributed on the surface of the bacteria, demonstrating homogeneous coating of the carrier with the recombinant enzyme. No fluorescence was detected on control inactivated *B. subtilis* incubated without the fusion protein.

### 3.3. Coated bacterial enzyme kinetics parameters

The kinetic constants of CBEs were determined for the hydrolysis of lactose and ONPG following Michaelis-Menten enzyme kinetics (Figure S1). The k_cat_ values were calculated based on the experimentally determined V_max_ values through non-linear regression, and using a molecular mass of 98 kDa for the catalytically active subunit. The V_max_ for ONPG is 9459.7 ± 504.9 U/mg, and the K_m_ for ONPG is 5.90 ± 0.77 mM. The V_max_ for lactose is 28.8 ± 2.3 U/mg, and the Km for lactose is 106.97 ± 15.91 mM. These K_m_ values are consistent with previously reported data.

Based on these parameters, it can be concluded that the obtained β-galactosidase exhibits a higher affinity towards ONPG compared to lactose. However, it is important to highlight that the K_m_ values obtained for lactose hydrolysis are significantly lower compared to other β-galactosidases. These results suggest that the obtained enzyme may have a significant application in the hydrolysis reaction of lactose to achieve low lactose concentrations.

**Table 1.**
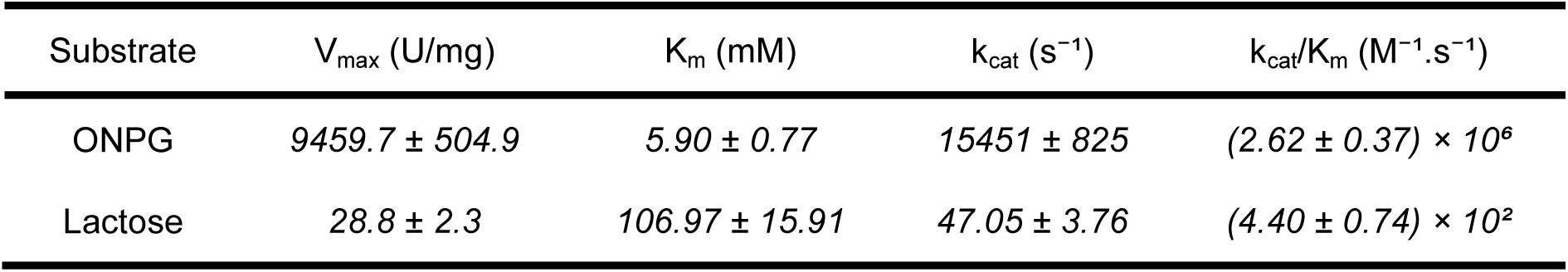
Kinetic parameters of CBEs for ONPG and lactose.

### 3.4. Effects of pH and temperature on enzyme activity

The influence of pH and temperature on the activity of free and immobilized β-galactosidase (CBE) was evaluated using both ONPG and lactose as substrates. Figure 3 shows the relative activity profiles under different conditions. Effect of pH (Figure 3A and 3B): For the free enzyme, maximum activity was observed at pH 6.0 with ONPG (Figure 3A) and at pH 6.0 with lactose (Figure 3B). Activity declined sharply when the pH moved away from these optima, particularly under acidic (pH < 5) or alkaline (pH > 7.5) conditions. In contrast, the immobilized enzyme exhibited a broader pH-activity profile, retaining more than 70% of its maximal activity between pH 5.5 and 7.5 for both substrates. The optimal pH remained at 6.0 for both substrates, but the immobilization conferred greater stability at extreme pH values.

**Figure 3.**
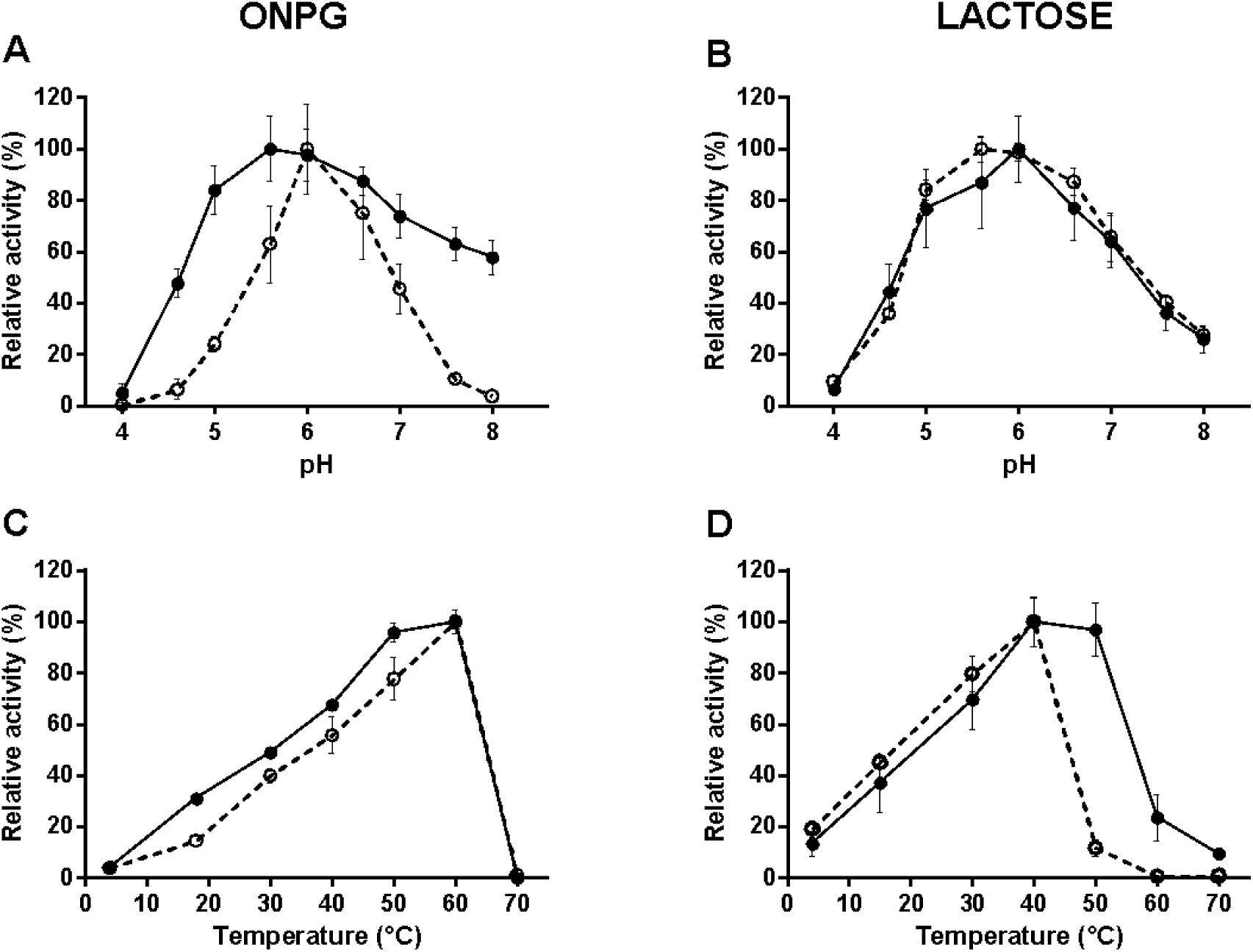
pH and temperature effect on the catalytic activity. Influence of pH (A and B) and temperature (C and D) in free enzyme (white circles) and CBEs (black circles) using ONPG (A and C) and lactose (B and D) as substrates. Values are presented as means and standard deviation of the means (*n*=3).

Effect of temperature (Figure 3C and 3D): The free enzyme showed maximum activity at 60 °C with ONPG (Fig. 3C) and at 40 °C with lactose (Fig. 3D), with activity declining rapidly above each optimum, likely due to thermal denaturation. For the immobilized enzyme, the optimal temperatures were similar (60 °C for ONPG, 50 °C for lactose), but the immobilized preparation retained significantly higher activity at elevated temperatures. For instance, at 50 °C with lactose as substrate, the CBE retained more than 90% of its maximal activity, whereas the free enzyme retained less than 10%. The immobilized enzyme also exhibited a broader operational temperature range, maintaining over 50% activity between 25 °C and 55 °C for both substrates. These results indicate that immobilization on the bacterial surface enhances enzyme thermostability and broadens the effective temperature range without substantially shifting the optimum values.

### 3.5. Thermal stability on enzyme activity

The thermal stability was investigated for CBEs and free BbgII-SlpA. To this purpose, we pre-incubated both enzyme forms at 45 °C for up to 390 min and measured residual activity at regular intervals using either ONPG or lactose as substrate (Fig. 4). When assayed with ONPG (Fig. 4A), the free enzyme lost activity rapidly during the first 60 min, retaining only approximately 20% of its initial activity after 390 min. In contrast, the immobilized CBE preparation maintained about 80% of its original activity under the same conditions, demonstrating a marked improvement in thermostability. A similar trend was observed when lactose was used as substrate (Fig. 4B). Again, the free enzyme showed a steep decline in activity, while the immobilized enzyme exhibited significantly higher residual activity throughout the entire incubation period. After 390 min at 45 °C, the free enzyme retained roughly 20% of its initial lactose hydrolysis activity, whereas the CBE retained approximately 80%. The stability profiles for both substrates, indicating that the protective effect of immobilization is independent of the substrate used.

**Figure 4.**
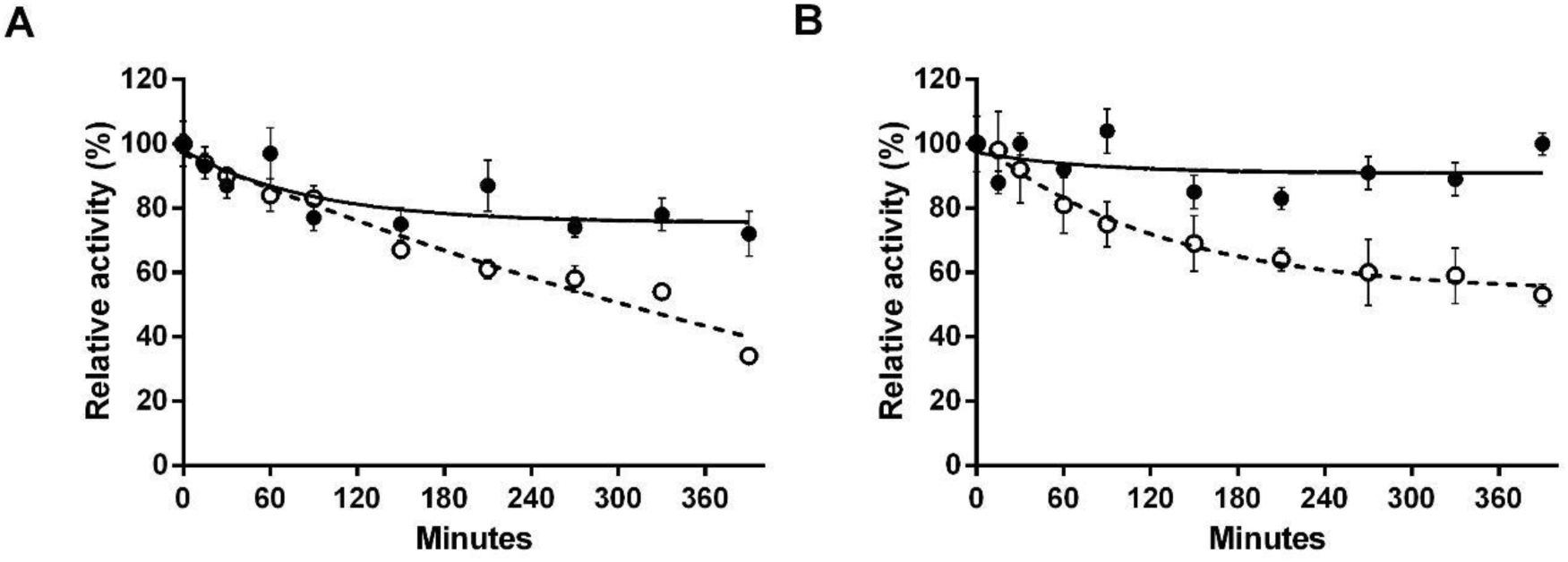
Thermal stability of free and immobilized BbgII-SlpA. The enzymes were pre-incubated at 45 °C for up to 390 min, and residual activity was measured at the indicated times using (A) ONPG or (B) lactose as substrate. CBEs (black circles); free enzyme (white circles). Values are mean ± SD (*n*=3). Residual activity is expressed as a percentage of the initial activity.

### 3.6. Cations effect on enzyme activity

The influence of various metal cations (Na^+^, K^+^, Mg^2+^, Mn^2+^, Ca^2+^, Ba^2+^, Cu^2+^, Zn^2+^, Co^2+^, Fe^2+^) on the activity of the immobilized CBEs was tested at three concentrations (1, 10, and 100 mM) using lactose as substrate. For comparison, the same cations were tested on the soluble enzyme BbgII-SlpA (Table 2). The results show that monovalent cations (Na^+^, K^+^) had little inhibitory effect on either enzyme form at all concentrations tested. Notably, K^+^ stimulated the soluble BbgII-SlpA at 100 mM (128% activity), while the immobilized CBEs was not significantly affected (92%). Divalent cations showed concentration-dependent inhibition. At 100 mM, Ca^2+^ completely abolished the activity of the soluble enzyme (0%) but the immobilized enzyme retained 36% activity. Cu^2+^ and Zn^2+^ also strongly reduced activity at high concentrations, with the immobilized enzyme showing higher residual activity (15% for Zn^2+^ vs. 0% for the free enzyme). Mn^2+^ was particularly detrimental to the soluble BbgII-SlpA at 100 mM (only 8% activity), while the immobilized CBEs retained 5% a similar level, not a protective effect. For Fe^2+^ at 100 mM, the soluble enzyme had only 5% activity, whereas the immobilized enzyme retained 74% activity, indicating a strong protective effect of immobilization against iron-induced inhibition. Overall, the immobilized enzyme (CBEs) exhibited similar cation sensitivity patterns to the soluble BbgII-SlpA, but was generally more tolerant to high concentrations of Ca^2+^, Zn^2+^, and especially Fe^2+^.

**Table 2.** Effect of different cations on the relative activity of immobilized CBEs and soluble BbgII-SlpA.

| Cation | BbgII-SlpA (% activity) |  |  | CBEs (% activity) |  |  |
| --- | --- | --- | --- | --- | --- | --- |
|  | 1 mM | 10 mM | 100 mM | 1 mM | 10 mM | 100 mM |
| $\text{Na}^+$ | 109 ± 14 | 107 ± 18 | 108 ± 17 | 94 ± 11 | 92 ± 14 | 92 ± 13 |
| $\text{K}^+$ | 96 ± 14 | 99 ± 10 | 128 ± 11 | 90 ± 11 | 99 ± 8 | 91 ± 8 |
| $\text{Mg}^{2+}$ | 103 ± 3 | 96 ± 11 | 93 ± 4 | 92 ± 3 | 91 ± 9 | 56 ± 3 |
| $\text{Mn}^{2+}$ | 105 ± 16 | 93 ± 6 | 8 ± 5 | 112 ± 13 | 97 ± 5 | 7 ± 4 |
| $\text{Ca}^{2+}$ | 112 ± 12 | 100 ± 5 | - | 105 ± 10 | 90 ± 4 | 36 ± 4 |
| $\text{Cu}^{2+}$ | 92 ± 5 | 43 ± 2 | - | 97 ± 4 | 13 ± 1 | - |
| $\text{Zn}^{2+}$ | 83 ± 5 | 80 ± 11 | - | 101 ± 4 | 67 ± 9 | 16 ± 5 |
| $\text{Co}^{2+}$ | 85 ± 5 | 83 ± 10 | 12 ± 5 | 90 ± 4 | 88 ± 7 | 6 ± 4 |
| $\text{Fe}^{2+}$ | 96 ± 6 | 63 ± 4 | 5 ± 4 | 101 ± 5 | 95 ± 3 | 73 ± 15 |

## 4. Discussion

One of the most widely used approaches in the production of lactose-reduced milk is the enzymatic hydrolysis of lactose using β-galactosidase. Enzymatic lactose hydrolysis requires high-purity β-galactosidase to avoid contaminating proteases that cause off-flavors [7][33]. Conventional multi-step purification is time-consuming, expensive, and low-yielding [15,16]. One-step affinity tags (e.g., polyhistidine, cellulose-binding domains) have been explored [17,20,21], but still require costly affinity matrices or specialized supports. In this work, we have developed a novel biocatalyst platform termed Coated Bacterial Enzymes (CBEs). This approach employs inactivated *B. subtilis* as a low-cost biological support to simultaneously purify and immobilize a recombinant β-galactosidase fused to the SlpA cell wall-binding domain, thereby expanding the scope of the SlpA-based tagging technology toward an enzymatic application.

Our results demonstrate that the chimeric enzyme BbgII-SlpA, expressed in *E. coli*, can be captured directly from crude lysate onto the surface of inactivated *B. subtilis* through specific affinity interaction, thereby forming the CBEs biocatalyst. The value obtained for the apparent dissociation constant (K_d_ = 16.2 μM) and maximum binding capacity (B_max_ = 144 μmol/g) indicate a stable and specific interaction, comparable to the K_d_ values previously reported for the SlpA domain with *Bacillus* surfaces [22,28,32]. The SlpA binding interaction with the cell wall of *B. subtilis* is mediated by recognition of teichoic and lipoteichoic acids (LTA) [34,35]. Since LTA is uniformly distributed across the Gram-positive cell wall, it is reasonable to expect a homogeneous distribution of the bound enzyme on the bacterial surface. This was confirmed by immunofluorescence microscopy (Fig. 2C), which showed a uniform green signal of BbgII-SlpA surrounding the entire carrier. Although previous studies have immobilized β-galactosidases on live bacteria using peptidoglycan-binding motifs such as LysM [36], or via surface display systems [37], those approaches rely on recombinant microorganisms that must maintain enzyme expression while alive. This introduces metabolic burden and regulatory concerns regarding genetically modified organisms. In contrast, our CBEs platform uses chemically inactivated *B. subtilis* as a non-recombinant carrier, decoupling enzyme production from the final biocatalyst and avoiding GMO-related limitations. However, the use of glutaraldehyde for inactivation, while effective, is not suitable for food-grade applications. Future work should explore alternative cross-linkers such as genipin, which is less toxic and food-compatible, to enhance the safety and regulatory acceptance of the CBE platform for dairy processing.

Subsequently, we characterized the catalytic properties of the CBE. The kinetic parameters obtained for the immobilized enzyme were compared with those of the recombinant BbgII from *Bifidobacterium bifidum* previously characterized by Goulas et al. [11]. For the synthetic substrate ONPG, our CBEs exhibited a slightly higher K_m_ (5.90 vs. 3.45 mM) but a substantially higher turnover number (k_cat_ = 15,451 s^−1^ vs. 4,041 s^−1^), resulting in a catalytic efficiency (k_cat_/K_m_) more than double that of the literature enzyme (2.62 × 10^6^ vs. 1.17 × 10^6^ M^−1^·s^−1^). For lactose, our CBEs showed a higher K_m_ (107 vs. 47 mM) and a slightly lower k_cat_ (47 vs. 70 s^−1^), leading to similar catalytic efficiencies (4.40 × 10^2^ vs. 1.48 × 10^3^ M^−1^·s^−1^). These comparisons highlight that the immobilized enzyme retains a remarkable turnover for the synthetic substrate ONPG while maintaining reasonable activity on lactose. The observed increase in the apparent Km for both substrates has been previously reported for β-galactosidases and can be attributed to mass transfer limitations and diffusional resistance [38,39]. In the case of our CBEs, this phenomenon could be associated with the microenvironment of the Gram-positive cell wall matrix, which may restrict substrate accessibility to the active site compared to the free enzyme. But further biophysical characterization would be required to confirm this hypothesis.

The effect of pH and temperature on the CBEs was evaluated in direct comparison with the free BbgII-SlpA. Both enzyme forms exhibited optimal activity at pH 6.0, consistent with the neutral lactase activity of BbgII from *B. bifidum* [11]. However, the immobilized preparation displayed a broader pH-activity profile using ONPG, retaining more than 60% of maximal activity between pH 5.5 and 8.0, whereas the free enzyme showed a sharper decline outside this optimum. This observation is consistent with oriented immobilization strategies seen in other recombinant systems [20,21,38]. While the free β-galactosidase from *B. bifidum* is often highly sensitive to pH fluctuations outside its narrow optimum, it has been previously reported that the cell-bound form benefits from changes in the microenvironment’s dielectric constant generated upon binding to the support [11,20]. Notably, BbgIII another β-galactosidase from *B. bifidum* possesses a multidomain membrane anchor motifs commonly found in cell wall-bound enzymes [40], and exhibits optimal activity in a broader neutral pH range (pH 6.4–6.8) compared to the more acidic optimum of BbgII (pH 5.4–5.8) [11]. This suggests that the native cell-wall association may contribute to a more robust pH profile, a feature that our SlpA-based immobilization strategy successfully recapitulates. Furthermore, this broader range is particularly advantageous for processing acidic dairy products such as whey or yogurt, where standard neutral lactases often lose significant activity [5]. Nevertheless, the pH stability of CBEs was evaluated only after short-term exposure; long-term stability at different pH values, particularly under industrial processing conditions, remains to be investigated.

Likewise, the optimal temperature for the immobilized enzyme was similar to that reported for free BbgII, with CBEs showing maximum activity at 60 °C for ONPG and 50 °C for lactose, compared to 50 °C and 40 °C, respectively, reported by Goulas et al. for the free BbgII isoenzyme [11]. The CBE exhibited markedly higher activity at elevated temperatures; for instance, at 50 °C using lactose, the immobilized enzyme retained >80% of its maximal activity compared to less than 10% for the free enzyme. The marked increase in thermal stability at 50 °C highlights the protective role of the SlpA anchoring to the cell wall. In free form, β-galactosidases are highly susceptible to thermal inactivation; for instance, some recombinant forms lose nearly all activity after short exposures to temperatures above 45–50 °C [10,41]. Immobilization on a rigid matrix acts as a form of molecular confinement, restricting the conformational mobility of the peptide chain and thereby preventing thermal denaturation, which preserves the native structure essential for catalytic activity [20]. Similar oriented immobilization techniques, such as using cellulose or chitin binding domains, have been shown to increase thermal stability compared to the free enzyme [42–44].

Regarding thermal stability, the immobilized CBE preparation exhibited a marked improvement compared to the free enzyme. When pre-incubated at 45 °C for up to 390 min, the free enzyme lost activity rapidly, retaining only approximately 20% of its initial activity for both ONPG and lactose hydrolysis. In contrast, the CBE maintained about 80% of its original activity under identical conditions, demonstrating a significant protective effect conferred by immobilization. Notably, the stability profiles for both substrates were nearly superimposable for each enzyme form, indicating that the protective effect is independent of the substrate used. This enhanced thermal stability is likely due to the rigid nature of the bacterial cell wall acting as a support matrix, which restricts conformational flexibility of the enzyme and prevents aggregation or unfolding at elevated temperatures [20,41]. Similar improvements in thermostability have been reported for other enzymes immobilized on cell-based supports or through oriented affinity tags, where the covalent or affinity binding reduces the entropy of the unfolded state, thereby shifting the thermal denaturation equilibrium toward the folded, active conformation [42–44]. In the case of BbgII, Goulas et al. reported that the free isoenzyme was highly susceptible to thermal inactivation, retaining only approximately 10% of its activity after 3 h at 50 °C [11]. Our results show that the CBE strategy not only preserves but significantly enhances the thermal resilience of BbgII, approaching the stability levels reported for the more thermostable BbgIV isoenzyme [11]. This improvement in operational stability is particularly relevant for industrial applications where prolonged processing times or moderate heating steps are involved. However, our thermal stability analysis was performed in a buffer solution rather than in a complex matrix such as milk, where other components (proteins, fats, salts) could influence enzyme stability. Future studies should evaluate CBE performance under real dairy processing conditions.

In addition to its enhanced thermal stability, the CBE platform also confers improved tolerance to cations that are naturally present in milk and can inhibit β-galactosidase activity. Our results showed that at 100 mM Ca^2+^, the free BbgII-SlpA lost all activity, whereas the CBE retained 36% of its activity. Similarly, Fe^2+^ at 100 mM reduced free enzyme activity to 6%, but the CBE retained 73% activity (Table 2). This protective effect is particularly relevant because milk contains significant concentrations of Ca^2+^ (approximately 30 mM) and other divalent cations [12]. The ability to withstand these inhibitory ions suggests that the CBE could be used directly in milk without prior demineralization or chelation steps, which would simplify the industrial process. Similar protective effects have been reported for other enzymes immobilized on solid supports, where the matrix acts as a physical barrier that shields the enzyme from inhibitory cations [45]. Together, the enhanced thermal stability and cation tolerance make the CBE platform a robust and practical alternative for lactose hydrolysis in dairy processing.

Finally, while the CBE platform demonstrates clear advantages in terms of purification and immobilization efficiency, several aspects require further investigation. These include the scalability of the carrier preparation, the potential for enzyme leakage during prolonged use, the stability of the biocatalyst under storage conditions, and the economic feasibility compared to existing commercial products. Addressing these aspects will be essential for translating the CBE technology from laboratory proof-of-concept to industrial implementation.

## 5. Conclusion

In this work, we have developed a novel biocatalyst platform termed Coated Bacterial Enzymes (CBEs), which employs chemically inactivated *Bacillus subtilis* as a low-cost biological support for the simultaneous purification and immobilization of recombinant proteins fused to the SlpA cell wall-binding domain. Using the dairy industry model enzyme BbgII from *Bifidobacterium bifidum*, we demonstrated that the SlpA-based anchoring system enables efficient one-step capture of the enzyme directly from crude *E. coli* lysate, with high binding capacity (B_max_ = 144 μmol/g) and specific affinity (K_d_ = 16.2 μM), comparable to other S-layer protein interactions. The resulting CBEs biocatalyst retained catalytic activity against both synthetic (ONPG) and natural (lactose) substrates, with kinetic parameters consistent with those reported for the native BbgII enzyme.

The immobilized enzyme exhibited enhanced operational robustness compared to its free counterpart, including broader pH and temperature activity ranges, significantly improved thermal stability (retaining ~80% activity after 390 min at 45 °C vs. ~20% for the free enzyme), and greater tolerance to inhibitory cations such as Ca^2+^ and Fe^2+^, which are naturally present in milk. These improvements are attributed to the rigid bacterial cell wall acting as a protective matrix that restricts conformational flexibility and reduces the accessibility of inhibitory ions to the active site. Notably, the CBEs platform decouples enzyme production from the final biocatalyst, avoiding the need for genetically modified organisms in the final product and overcoming regulatory constraints associated with live bacterial carriers.

While our results highlight the potential of CBEs for industrial lactose hydrolysis, further studies are needed to evaluate the reusability of the biocatalyst in continuous processes and its performance in complex matrices such as skim milk or whey. Additionally, optimization of the antigen-binding capacity and exploration of other industrially relevant enzymes could further expand the versatility of this platform.

In conclusion, the CBE platform offers a straightforward, cost-effective, and robust alternative for obtaining high-purity, immobilized β-galactosidase, with promising applications in the production of lactose-reduced milk and other dairy products. This work successfully expands the scope of the SlpA-based tagging technology from protein purification to functional enzyme immobilization, paving the way for its broader adoption in industrial biotechnology.

## Authors contribution statement

RAC and HI performed the experimental work, generated the figures, and contributed to the writing and discussion of the manuscript. HMS conducted the cations experiments and contributed to the writing. SAE generated the script for the determination of enzyme kinetics. CSF participated in the writing and discussion of the manuscript’s results. OGE, as the principal investigator and project director, contributed to the experimental design, interpretation of the results, and writing of the manuscript. All authors contributed to the development of the article and approved the final submitted version.

## Supplementary Data

Ramírez Gutiérrez, A. C., Harguindeguy, I., Homse, S. M., Alan Eric, S., Cavalitto, S. F., & Ortiz, G. (2026). Coated Bacterial Enzymes: A one-step approach for enzymatic purification and immobilization. Zenodo. https://doi.org/10.5281/zenodo.20934760

