## Supplementary figures and images for "Coated Bacterial Enzymes: A one-step approach for enzymatic purification and immobilization"

### Figure S1

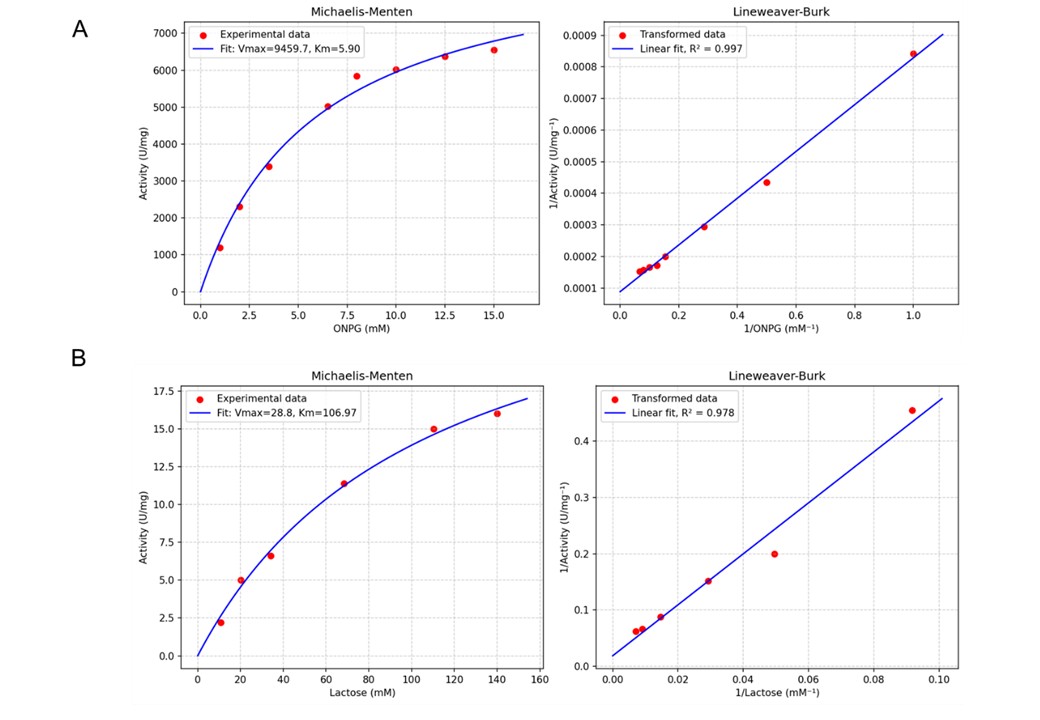
